# CRISPR-Cas12a target binding unleashes single-stranded DNase activity

**DOI:** 10.1101/226993

**Authors:** Janice S. Chen, Enbo Ma, Lucas B. Harrington, Xinran Tian, Jennifer A. Doudna

## Abstract

CRISPR-Cas12a (Cpf1) proteins are RNA-guided DNA targeting enzymes that bind and cut DNA as components of bacterial adaptive immune systems. Like CRISPR-Cas9, Cas12a can be used as a powerful genome editing tool based on its ability to induce genetic changes in cells at sites of double-stranded DNA (dsDNA) cuts. Here we show that RNA-guided DNA binding unleashes robust, non-specific single-stranded DNA (ssDNA) cleavage activity in Cas12a sufficient to completely degrade both linear and circular ssDNA molecules within minutes. This activity, catalyzed by the same active site responsible for site-specific dsDNA cutting, indiscriminately shreds ssDNA with rapid multiple-turnover cleavage kinetics. Activation of ssDNA cutting requires faithful recognition of a DNA target sequence matching the 20-nucleotide guide RNA sequence with specificity sufficient to distinguish between closely related viral serotypes. We find that target-dependent ssDNA degradation, not observed for CRISPR-Cas9 enzymes, is a fundamental property of type V CRISPR-Cas12 proteins, revealing a fascinating parallel with the RNA-triggered general RNase activity of the type VI CRISPR-Cas13 enzymes.

**One Sentence Summary:** Cas12a (Cpf1) and related type V CRISPR interference proteins possess non-specific, single-stranded DNase activity upon activation by guide RNA-dependent DNA binding.

## Main Text

CRISPR-Cas adaptive immunity in bacteria and archaea uses RNA-guided nucleases to target and degrade foreign nucleic acids (*1, 2*). The CRISPR-Cas9 family of proteins has been widely deployed for gene editing applications (*3, 4*) based on the precision of double-stranded DNA (dsDNA) cleavage induced by two catalytic domains, RuvC and HNH, at sequences complementary to a guide RNA sequence (*5, 6*). A second family of enzymes, CRISPR-Cas12a (Cpf1), uses a single RuvC catalytic domain for guide RNA-directed dsDNA cleavage (*7-12*) (**Fig. 1a**). Distinct from Cas9, Cas12a enzymes recognize a T-rich protospacer adjacent motif (PAM) (*7*), catalyze their own guide RNA (crRNA) maturation (*13*) and generate a PAM-distal dsDNA break with staggered 5’ and 3’ ends (*7*), features that have attracted interest for gene editing applications (*14-16*). However, the substrate specificity and DNA cleavage mechanism of Cas12a are yet to be fully elucidated.

**Fig. 1.**
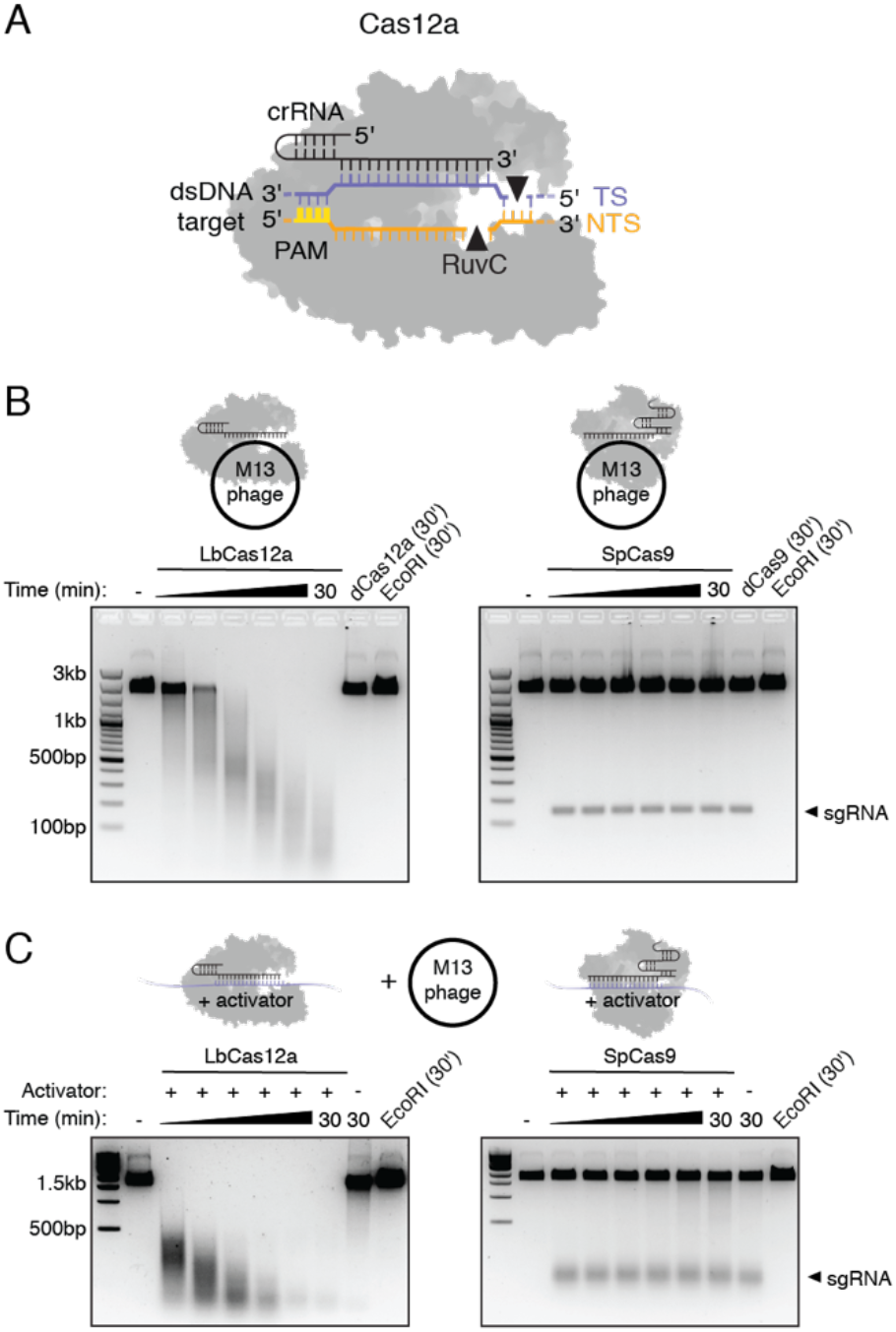
Cas12a target recognition activates non-specific single-stranded DNA cleavage. (**A**) Cas12a-crRNA complex binds a dsDNA substrate and generates a 5’ overhang staggered cut using a single RuvC nuclease. (**B, C**) Representative M13 ssDNA cleavage timecourses with purified LbCas12a (left) and SpCas9 (right) complexed with a (**B**) guide RNA complementary to M13 phage or (**C**) a guide RNA and complementary ssDNA activator with no sequence homology to M13 phage.

While investigating substrate requirements for Cas12a activation, we tested *Lachnospiraceae* bacterium ND2006 Cas12a (LbCas12a) for guide RNA-directed singlestranded DNA (ssDNA) cleavage, a capability of diverse CRISPR-Cas9 orthologs (*17, 18*). Purified LbCas12a or *Streptococcus pyogenes* Cas9 (SpCas9) proteins (**fig. S1**) were assembled with guide RNA sequences targeting a circular, single-stranded M13 DNA phage. In contrast to SpCas9, we were surprised to find that LbCas12a induced rapid and complete degradation of M13 by a cleavage mechanism that could not be explained by sequence-specific DNA cutting (**Fig. 1b**). This ssDNA shredding activity, not observed using an LbCas12a protein containing inactivating mutations in the RuvC catalytic domain, raised the possibility that a target-bound LbCas12a could degrade any ssDNA, regardless of complementarity to the guide RNA. To test this idea, we assembled LbCas12a or SpCas9 with a different guide RNA and complementary ssDNA with no sequence homology to M13 phage genome sequence, and added M13 DNA to the reaction. Remarkably, LbCas12a catalyzed M13 degradation only in the presence of this complementary ssDNA “activator”, an activity not observed for SpCas9 (**Fig. 1c**). These findings reveal that binding of LbCas12a to a guide-complementary ssDNA unleashes robust, non-specific ssDNA *trans-cleavage* activity.

We next investigated the requirements for LbCas12-catalyzed trans-cleavage activity. Using a fluorophore quencher (FQ)-labeled reporter assay (*19, 20*), we assembled LbCas12a with its crRNA and either a complementary ssDNA, dsDNA or single-stranded RNA (ssRNA), and introduced an unrelated ssDNA- or ssRNA-FQ reporter in *trans* (**fig. S2**). Both the crRNA-complementary ssDNA or dsDNA (the activator) triggered LbCas12a to cleave the ssDNA-FQ reporter substrate (**fig. S2A**). However, ssRNA was neither capable of activating trans-cleavage nor susceptible to degradation by LbCas12a (**fig. S2B**), confirming that LbCas12a harbors a DNA-activated general DNase activity.

To determine how LbCas12a-catalyzed ssDNA cleavage activity relates to site-specific dsDNA cutting, we tested the length requirements of the target strand (TS) and non-target strand (NTS) for LbCas12a activation using radiolabeled oligonucleotides. Although TS cutting occurred irrespective of the NTS length (**figs. S3A-B**), NTS cleavage occurred only when the TS contained at least 15 nucleotides (nt) of complementarity with the crRNA (**fig. S3C**). This showed that TS recognition is a prerequisite for NTS cutting. To test whether LbCas12a remains active for non-specific ssDNA cleavage after sequence-specific binding and cleavage of a dsDNA substrate, we first cut a dsDNA plasmid with an LbCas12a-crRNA complex, and introduced an unrelated dsDNA or ssDNA to the reaction (**Fig. 2a**). Whereas non-specific dsDNA substrate remained intact, the ssDNA was rapidly degraded in a RuvC-domain dependent manner (**Fig. 2a; figs. S4, S5**). Together, these results show that RNA-guided DNA binding activates LbCas12a for both site-specific dsDNA cutting and non-specific ssDNA *trans*-cleavage.

**Fig. 2.**
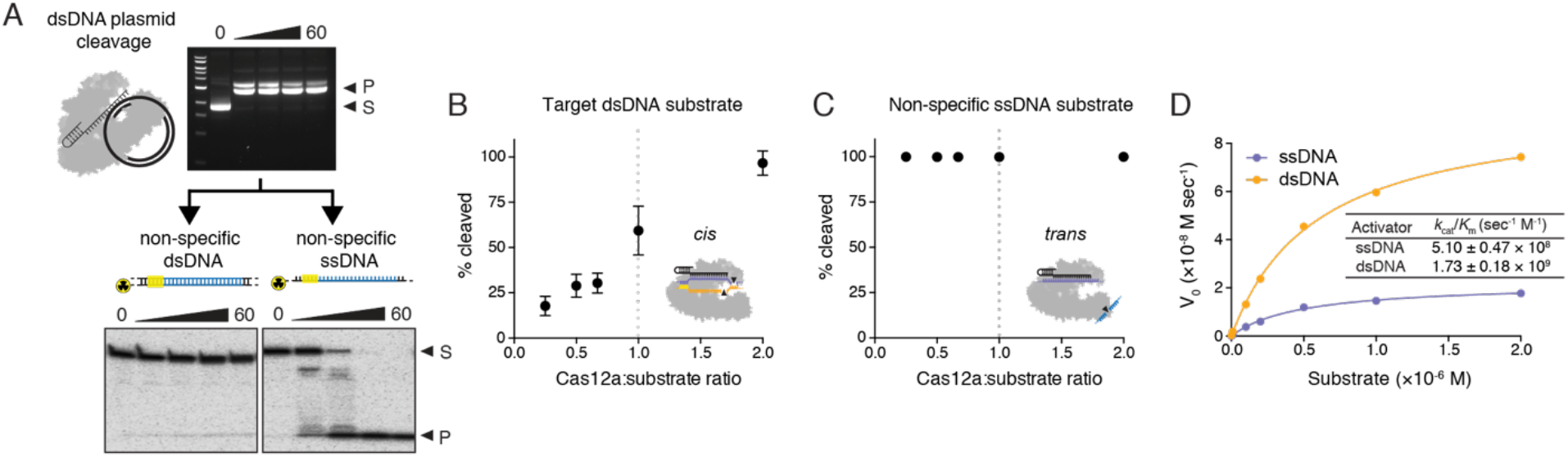
Kinetics of Cas12a ssDNA *trans*-cleavage. (**A**) Sequence-specific plasmid DNA cleavage reactions by LbCas12a-crRNA (top) were introduce to a separate radiolabeled dsDNA or ssDNA substrate of unrelated sequence (bottom); timecourses represent minutes. (**B**) Target dsDNA or (**C**) non-specific ssDNA incubated with molar ratios of LbCas12a-crRNA as indicated. Each point represents the mean quantified percent cleavage after 30 minutes at 37°C, at which time the reaction was at completion. Error bars represent the mean ± s.d., where *n* = 3. (**D**) Representative Michaelis-Menten plot for LbCas12a-catalyzed ssDNA trans-cleavage using a dsDNA or ssDNA activator. Measured *k_cat_/K_m_* values report mean ± s.d., where *n* = 3.

The rapid degradation of a *trans* substrate suggested that the kinetics of LbCas12a-catalyzed site-specific dsDNA (cis-) cleavage and non-specific ssDNA *(trans-)* cleavage are fundamentally different. Stoichiometric titration experiments showed that cis-cleavage is single-turnover (*21*) (**Fig. 2b**), whereas trans-cleavage is multiple-turnover (**Fig. 2c**). Using the FQ assay, we found that LbCas12a-crRNA bound to a ssDNA activator molecule catalyzed ssDNA trans-cleavage at a rate of ~250 turnovers per second and a catalytic efficiency (k_cat_/K_m_) of 5.1 ×10^8^ s^-1^ M^-1^. When bound to a dsDNA activator, LbCas12a-crRNA catalyzed ~1250 turnovers per second with a catalytic efficiency approaching the rate of diffusion (*22*) with a k_ca_t/K_m_ of 1.7×10^9^ s^-1^ M^-1^ (**Fig. 2d; fig. S6**). These differences suggest that the NTS of the dsDNA activator helps stabilize the Cas12a complex in an optimal conformation for *trans-* ssDNA cutting.

We next tested the specificity of trans-cleavage activation using either a ssDNA or dsDNA activator. We found that the PAM sequence required for dsDNA binding by CRISPR-Cas12a (*21*) is critical for catalytic activation by a crRNA-complementary dsDNA (*8, 23*), but not for a crRNA-complementary ssDNA (**Fig. 3a**). Two base-pair (bp) mismatches introduced along the crRNA-complementary sequence of either a ssDNA or dsDNA activator molecule slowed the trans-cleavage rate of a ssDNA-FQ reporter by up to ~100 fold, depending on the mismatch position. For only the dsDNA activator, alterations to the PAM sequence or mismatches between the crRNA and PAM-adjacent “seed region” also had large inhibitory effects on trans-ssDNA cleavage activity (**Fig. 3b; fig. S7**), similar to the mismatch tolerance pattern observed in Cas12a off-target studies (*24-26*). Together, these data are consistent with PAM-mediated dsDNA target binding and the role of base pairing between the crRNA and the target strand to activate trans-ssDNA cutting.

**Fig. 3.**
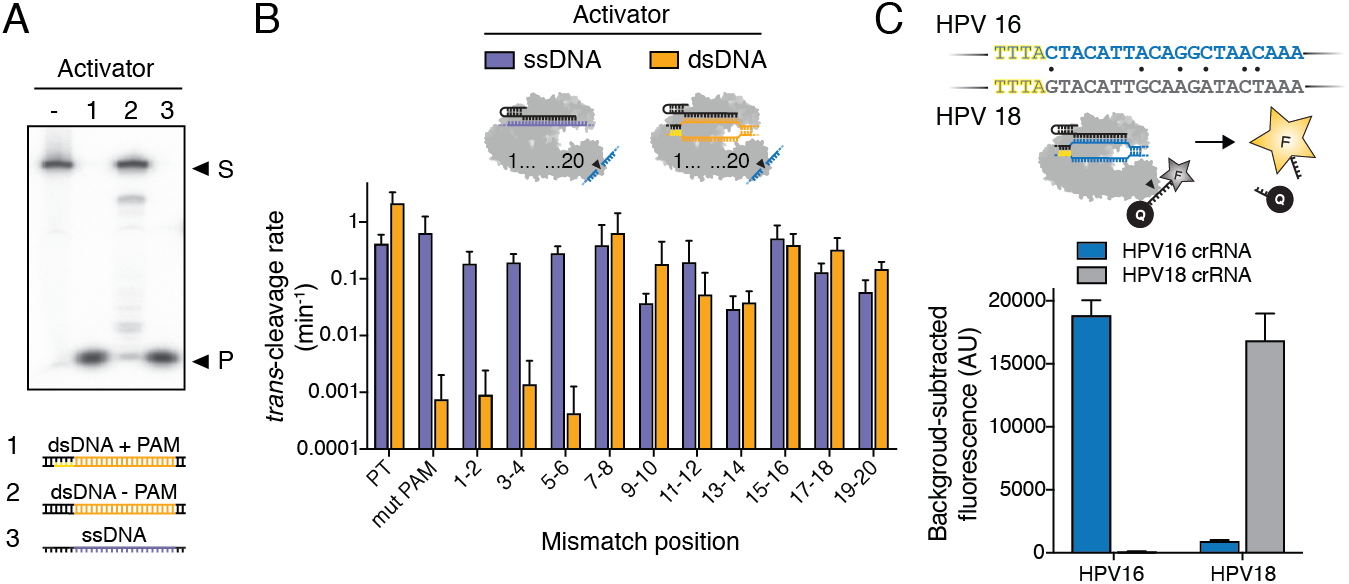
Specificity of *trans*-cleavage activation. (**A**) LbCas12a-crRNA in the absence or presence of indicated activator, incubated with a radiolabeled non-specific ssDNA substrate (S) for 30 min at 37°C; products (P) resolved by denaturing PAGE. (**B**) Observed trans-cleavage rates for LbCas12a using a ssDNA or dsDNA activator with indicated mismatches; rates represent the average of three different targets measured in triplicate, and error bars represent mean ± s.d., where *n* = 9. (**C**) 20 base-pair target sequence with adjacent TTTA PAM (yellow) in related dsDNA human papillomavirus (HPV) serotypes 16 (HPV16) and 18 (HPV18) differs by six base pairs (black dots). Binding to a cognate HPV sequence activates LbCas12a transcleavage of the DNaseAlert substrate (see **Methods**) and generates a fluorescent signal. Background-subtracted maximum fluorescence values represent mean ± s.d., where *n* = 3.

We next explored whether LbCas12a-catalyzed trans-ssDNA cleavage can distinguish between two closely-related dsDNA viruses, human papillomavirus (HPV) serotypes 16 (HPV16) and 18 (HPV18) (*27*). Within the HPV16 and HPV18 genomes, we selected a 20 nt target sequence located next to a TTTA PAM that varied by only six base pairs between the two serotypes (**Fig. 3c**). Plasmids containing a ~500 bp fragment of the HPV16 or HPV18 genome, including the target sequence, were incubated with the LbCas12a-crRNA complex targeting either the HPV16 or HPV18 fragment and a fluorescent ssDNA reporter. Robust LbCas12a-catalyzed ssDNA trans-cleavage occurred only in the presence of the cognate HPV target (**Fig. 3c; fig. S8**), suggesting that dsDNA recognition specificity and trans-cleavage activity could in principle be extended to detect any dsDNA sequence.

We wondered if this trans-ssDNA cutting activity might be a property of Cas12a, and perhaps more evolutionarily distinct type V CRISPR effector proteins, considering that all type V effectors contain a single RuvC nuclease domain (*28, 29*). We therefore purified Cas12a orthologs from *Acidaminococcus sp*. (AsCas12a) and *Francisella novicida* (FnCas12a), as well as a Cas12b protein from *Alicyclobacillus acidoterrestris* (AaCas12b) to test for *cis-* and *trans-ssDNA* cleavage activity. With varying efficiencies, all of these homologs catalyzed non-specific ssDNase cleavage when assembled with crRNA and a complementary ssDNA activator (**Fig. 4a**), suggesting that target-dependent activation of non-specific ssDNA cleavage is a fundamental feature of all type V CRISPR-Cas12-family proteins.

**Fig. 4.**
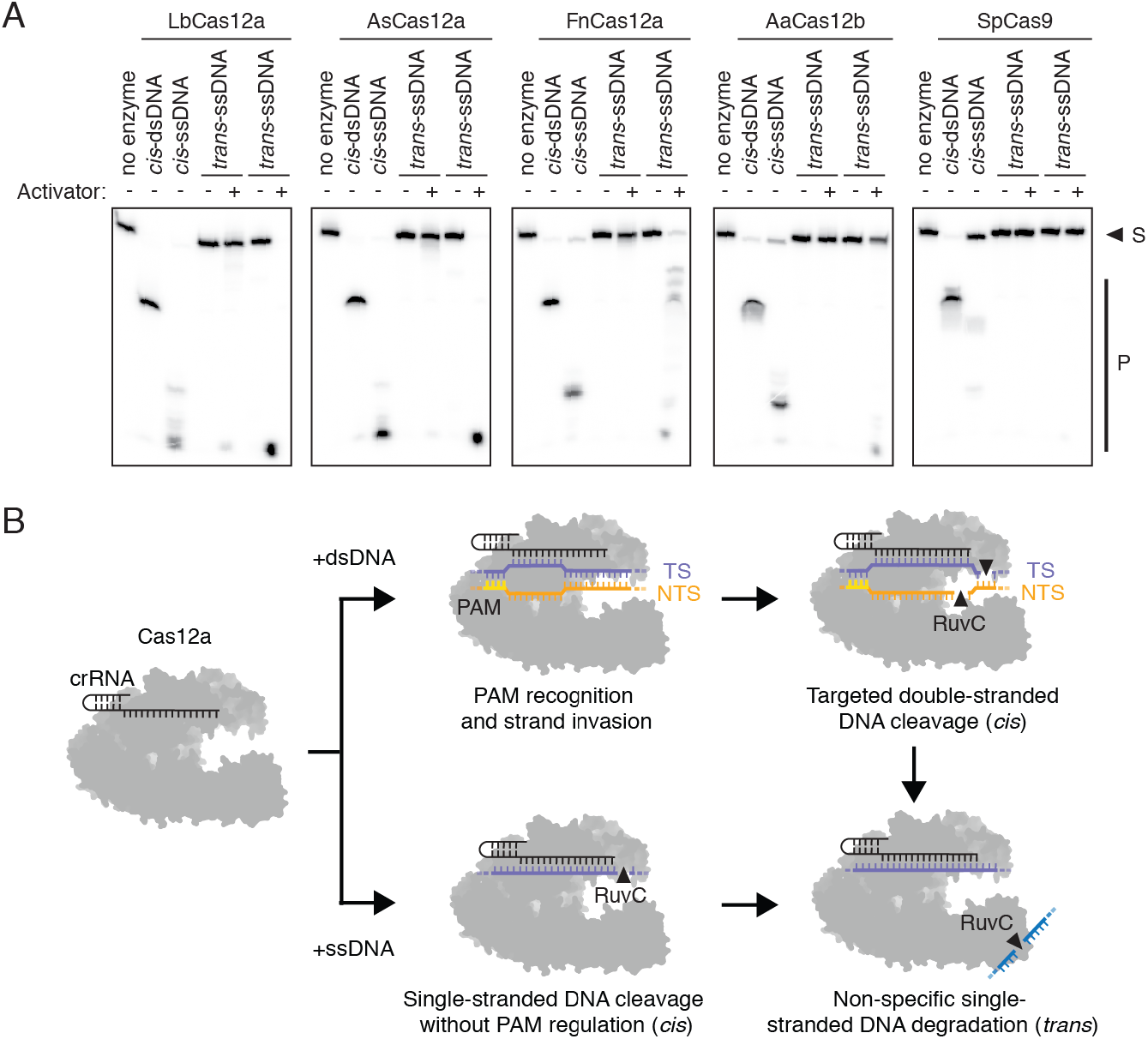
Activator-dependent, non-specific ssDNA cleavage activity is conserved across type V CRISPR systems. (**A**) Radiolabeled *cis* (complementary) or *trans* (non-complementary) substrates were incubated with Cas12-crRNA or Cas9-sgRNA in the presence or absence of a ssDNA activator for 30 min at 37°C (or 47.5°C for AaCas12b); substrate (S) and nucleotide products (P) resolved by denaturing PAGE. (**B**) Model for PAM-dependent and PAM-independent activation of *cis* and *trans*-cleavage by Cas12a.

Together, these findings support a unifying mechanism of target interference that begins with the Cas12a-guide RNA complex binding to a complementary DNA sequence in a PAM-dependent (dsDNA) or PAM-independent (ssDNA) manner (**Fig. 4b**). Within a host bacterium, such enzyme activation could provide simultaneous protection from both dsDNA and ssDNA phages, and could also target ssDNA sequences that arise temporarily during phage replication or transcription (*30*). For genome editing applications, this newly identified activity may also degrade single-stranded oligonucleotide donors that are often used for homology directed repair (*31*), altering DNA repair outcomes. Finally, these results reveal unexpected functional convergence of target-dependent trans-cleavage activity between the DNA-targeting type V and RNA-targeting type VI CRISPR effector enzymes (*19, 32, 33*) and broaden the antiviral capabilities of CRISPR interference.

## Acknowledgments

We thank O. Mavrothalassitis, D. Burstein, D. Lee and members of the Doudna laboratory for comments and discussions. This research was supported in part by the Allen Distinguished Investigator Program, through The Paul G. Allen Frontiers Group, and the National Science Foundation (MCB-1244557 to J.A.D.). J.S.C. and L.B.H. are supported by National Science Foundation Graduate Research Fellowships. J.A.D. is an Investigator of the Howard Hughes Medical Institute and executive director of the Innovative Genomics Institute at the University of California, Berkeley and the University of California, San Francisco. J.A.D. is a co-founder of Editas Medicine, Intellia Therapeutics, and Caribou Biosciences and a scientific adviser to Caribou, Intellia, eFFECTOR Therapeutics and Driver. The Regents of the University of California have patents pending for CRISPR technologies on which the authors are inventors.

## Materials and Methods

### Protein expression and purification

SpCas9 and Cas12 proteins and mutants were cloned into a custom pET-based expression vector containing an N-terminal 10×His-tag, maltose-binding protein (MBP) and TEV protease cleavage site. Point mutations were introduced by around-the-horn PCR and verified by DNA sequencing. Proteins were purified as described (*1*), with the following modifications: *E. coli* BL21(DE3) containing SpCas9 or Cas12 expression plasmids were grown in Terrific Broth at 16°C for 14 hr. Cells were harvested and resuspended in Lysis Buffer (50 mM Tris-HCl, pH 7.5, 500 mM NaCl, 5% (v/v) glycerol, 1 mM TCEP, 0.5 mM PMSF and 0.25 mg/ml lysozyme), disrupted by sonication, and purified using Ni-NTA resin. After overnight TEV cleavage at 4°C, proteins were purified over an MBPTrap HP column connected to a HiTrap Heparin HP column for cation exchange chromatography. The final gel filtration step (Superdex 200) was carried out in elution buffer containing 20 mM Tris-HCl, pH 7.5, 200 mM NaCl (or 250 mM NaCl for AaCas12b), 5% (v/v) glycerol and 1 mM TCEP. All proteins tested in this study are shown in **fig. S1**.

### Nucleic acid preparation

DNA substrates were synthesized commercially (IDT). For FQ-reporter assays, activator DNA duplexes were prepared by annealing 5-fold molar excess of the NTS to TS in 1 × hybridization buffer (20 nM Tris-Cl, pH 7.5, 100 mM KCl, 5 mM MgCl_2_), heating at 95°C and slow-cooling on the benchtop. HPV16 and HPV18 fragments were synthesized as gBlocks (IDT) and cloned into a custom pET-based vector via Gibson assembly. For radiolabeled cleavage assays, PAGE-purified DNA oligos were prepared as described.

sgRNA templates were PCR amplified from a pUC19 vector or overlapping primers containing a T7 promoter, 20 nucleotide target sequence and an sgRNA scaffold. The amplified PCR product served as the DNA template for *in vitro* transcription reactions, which were performed as described (*1*). crRNAs were transcribed *in vitro* using a single-stranded DNA template containing a T7 promoter, repeat and spacer in the reverse complement orientation, which was annealed to T7 forward primer in 1 × hybridization buffer. All DNA and RNA substrates are listed in **Table S1**.

### DNA cleavage assays

Generally, Cas12a-mediated cleavage assays were carried out in Cleavage Buffer consisting of 20 mM HEPES (pH 7.5), 150 mM KCl, 10 mM MgCl_2_, 1% glycerol and 0.5 mM DTT. For M13-targeting assays, 30 nM Cas12a was pre-assembled with either 36 nM of M13-targeting crRNA *(cis)* or with 36 nM of crRNA and 40 nM complementary ssDNA (activator) with no sequence homology to M13 *(trans)* at 37°C for 10 min. The reaction was initiated by adding 10 nM M13mp18 ssDNA (New England Biolabs) and incubated at 37°C for indicated timepoints. Reactions were quenched with DNA loading buffer (30% (v/v) glycerol, 0.25% (w/v) bromophenol blue, 0.25% (w/v) xylene cyanol) containing 15 mM EDTA and separated by 1.5% agarose gel pre-stained with SyberGold.

For radiolabeled cleavage assays, the substrates used were 5’-end-labeled with T4 PNK (NEB) in the presence of gamma ^32^P-ATP. For dsDNA substrates, the non-target strand was first 5’-end-labeled and then annealed with excess corresponding target strand. The concentrations of Cas12a (or SpCas9), guide RNA and ^32^P-labeled substrates used in the reaction were 30 nM, 36 nM and 1-3 nM, respectively. Reactions were incubated for 30 min (unless otherwise stated) at 37°C (or 47.5°C for the thermophilic AacCas12b) and quenched with formamide loading buffer (final concentration 45% formamide and 15 mM EDTA, with trace amount of xylene cyanol and bromophenol blue) for 3 min at 90°C. The substrates and products were resolved by 12% urea-denaturing PAGE gel and quantified with Amersham Typhoon (GE Healthcare).

For substrate turnover studies, the pre-assembled Cas12a-crRNA or Cas12a-crRNA-activator (target ssDNA or dsDNA) were incubated at 37°C for 10 min, and 30 nM of the preassembled RNP were used for each reaction with various substrate concentrations at 15, 30, 45, and 60 nM, respectively.

### Fluorophore quencher (FQ)-labeled reporter assays

LbCas12a-crRNA complexes were preassembled by incubating 200 nM LbCpf1 with 250 nM crRNA and 4 nM activator (ssDNA, dsDNA or ssRNA) at 37°C for 30 min. The reaction was initiated by diluting LbCas12a complexes to a final concentration of 50 nM LbCas12a: 62.5 nM crRNA: 1 nM activator in a solution containing 1× Binding Buffer (20 mM Tris-HCl, pH 7.5, 100 mM KCl, 5 mM MgCl_2_, 1 mM DTT, 5% glycerol, 50 μg ml^-1^ heparin) and 50 nM DNaseAlert substrate™ (IDT) or custom ssDNA/ssRNA FQ reporter substrates (**Table S1**). Reactions were incubated in a fluorescence plate reader (Tecan Infinite Pro F200) for up to 120 minutes at 37°C with fluorescence measurements taken every 30 seconds (DNaseAlert substrate = λex: 535 nm; λem: 595 nm, custom ssDNA/ssRNA FQ substrates = λex: 485 nm; λem: 535 nm).

For *trans*-cleavage rate determination, background-corrected fluorescence values were calculated by subtracting fluorescence values obtained from reactions carried out in the absence of target plasmid. The resulting data were fit to a single exponential decay curve (GraphPad Software), according to the following equation: Fraction cleaved = A × (1 – exp(-k × t)), where A is the amplitude of the curve, k is the first-order rate constant, and t is time.

For Michaelis-Menten analysis, LbCas12a-crRNA-activator (target ssDNA or dsDNA) complexes were prepared as described above, and reaction was initiated by diluting LbCas12a complexes to a final concentration of 5 nM LbCas12a: 6.25 nM crRNA: 0.1 nM activator (effective complex = 0.1 nM) in a solution containing 1× Binding Buffer and 0.001, 0.01, 0.1, 0. 2, 0.5, 1 or 2 uM of DNaseAlert™ substrate (IDT). Reactions were incubated in a fluorescence plate reader for up to 30 minutes at 37°C with fluorescence measurements taken every 30 seconds (λex: 535 nm; λem: 595 nm). The initial velocity (V_0_) was calculated by fitting to a linear regression and plotted against the substrate concentration to determine the Michaelis-Menten constants (GraphPad Software), according to the following equation: Y = (V_max_ × X)/(*K*_m_ + X), where X is the substrate concentration and Y is the enzyme velocity. The turnover number (k_cat_) was determined by the following equation: *k*_cat_ = V_max_/E_t_, where E_t_ = 0.1 nM.

HPV detection assays were performed as above, with the following modifications: LbCas12a was pre-assembled with an HPV16 or HPV18-targeting crRNA and diluted in a solution containing 1× Binding Buffer, 50 nM DNaseAlert substrate™ (IDT) and 1, 10, 100, or 1000 nM of HPV16- or HPV18-containing plasmids. Reactions were incubated in a fluorescence plate reader for 60 minutes at 37°C with fluorescence measurements taken every 30 seconds (λex: 535 nm; λem: 595 nm).

---

**Fig. S1.**
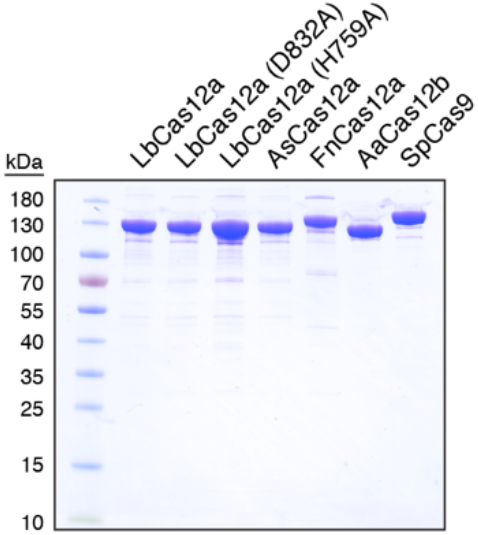
Purification of Cas12 and Cas9 proteins. SDS-PAGE gel of all purified Cas12 and Cas9 proteins used in this study.

**Fig. S2.**
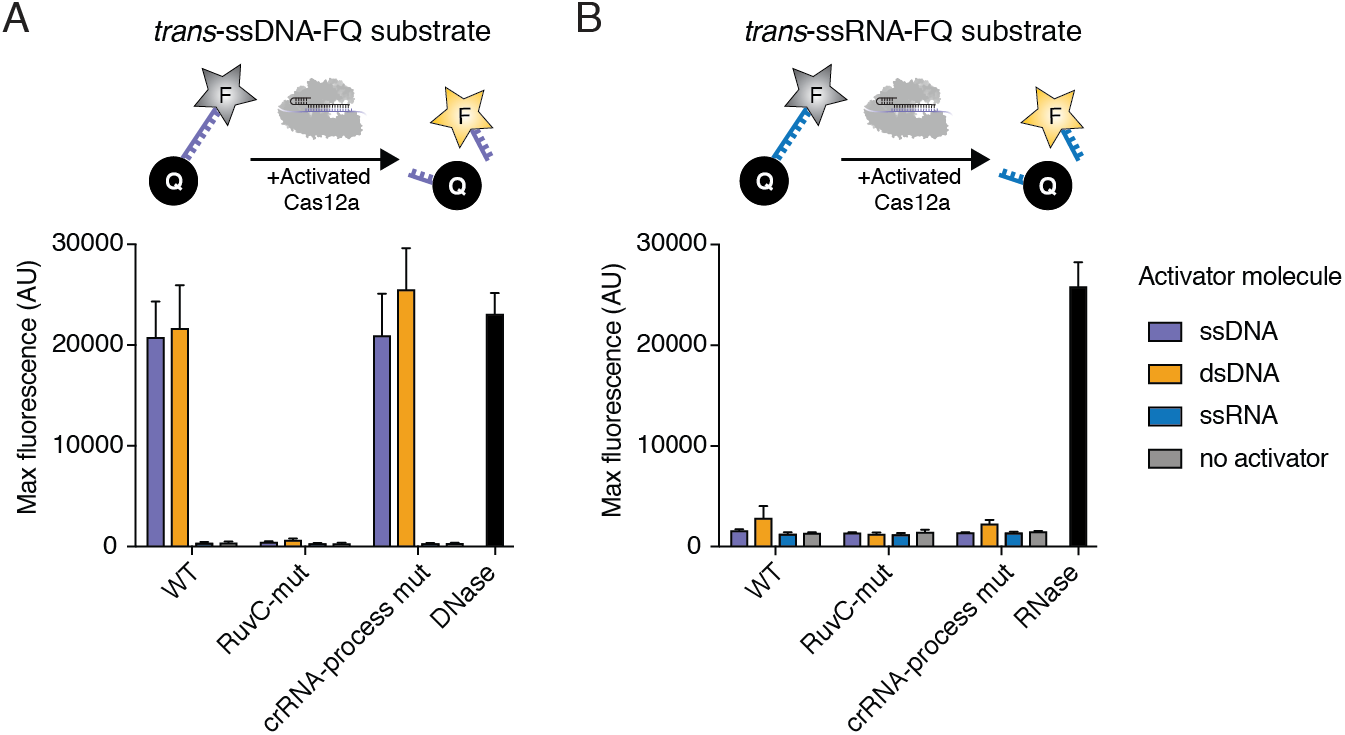
LbCas12a is a DNA-activated general DNase. Quantification of maximum fluorescence signal generated after incubating LbCas12a-crRNA-activator with a custom (**A**) trans-ssDNA-FQ or (**B**) trans-ssRNA-FQ reporter for 1h at 37°C, with DNase I or RNase A controls where indicated. Error bars represent the mean ± s.d., where *n* = 3.

**Fig. S3.**
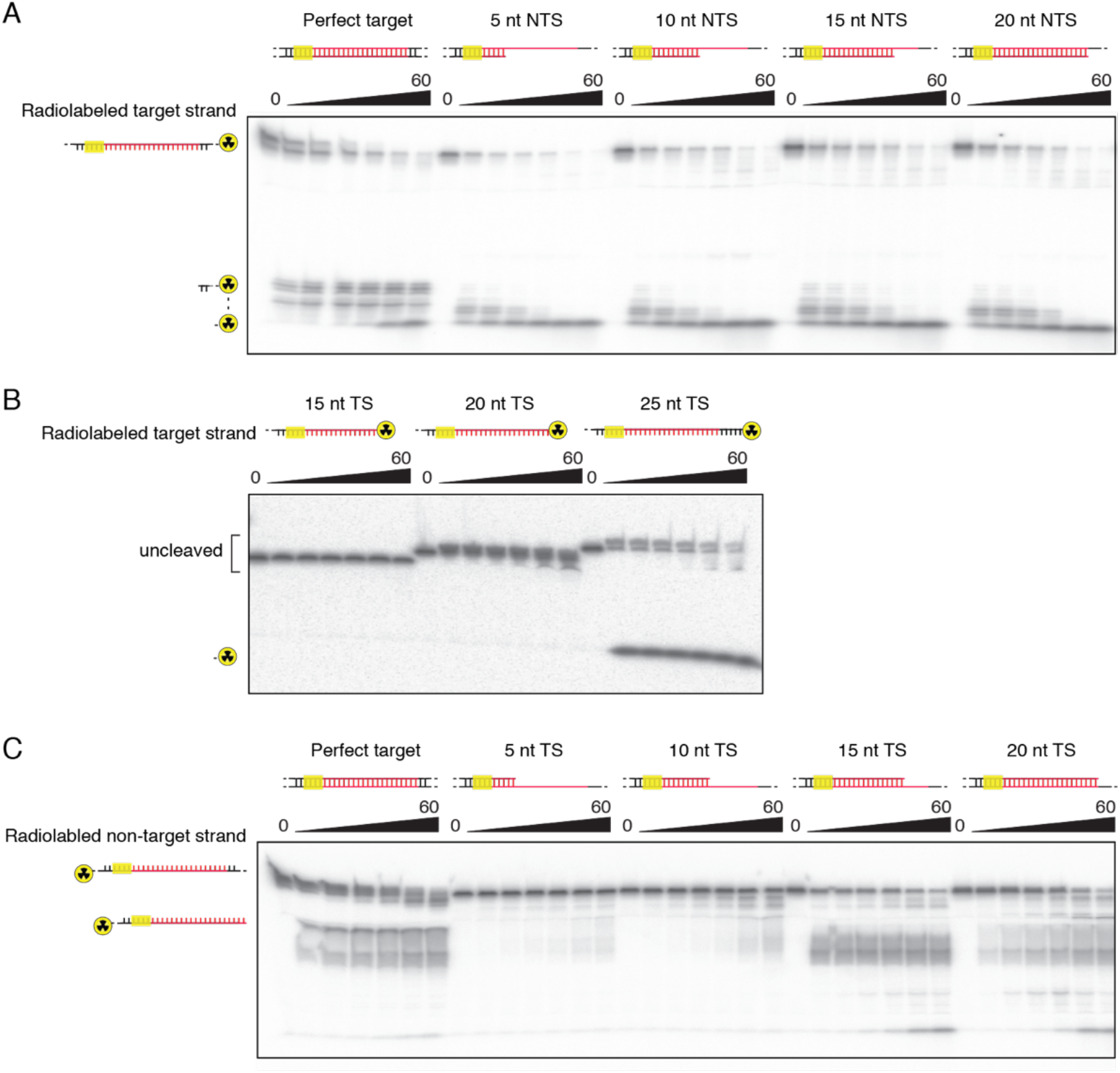
Target strand recognition is a pre-requisite for single-stranded DNA cleavage. Cleavage timecourse assays using LbCas12a with (**A**) truncated target strand (TS) annealed to a radiolabeled non-target strand (NTS), (**B**) truncated and radiolabeled TS only, and (**C**) truncated NTS annealed to a radiolabeled TS. Timecourses represent minutes and cleavage products are resolved by denaturing PAGE.

**Fig. S4.**
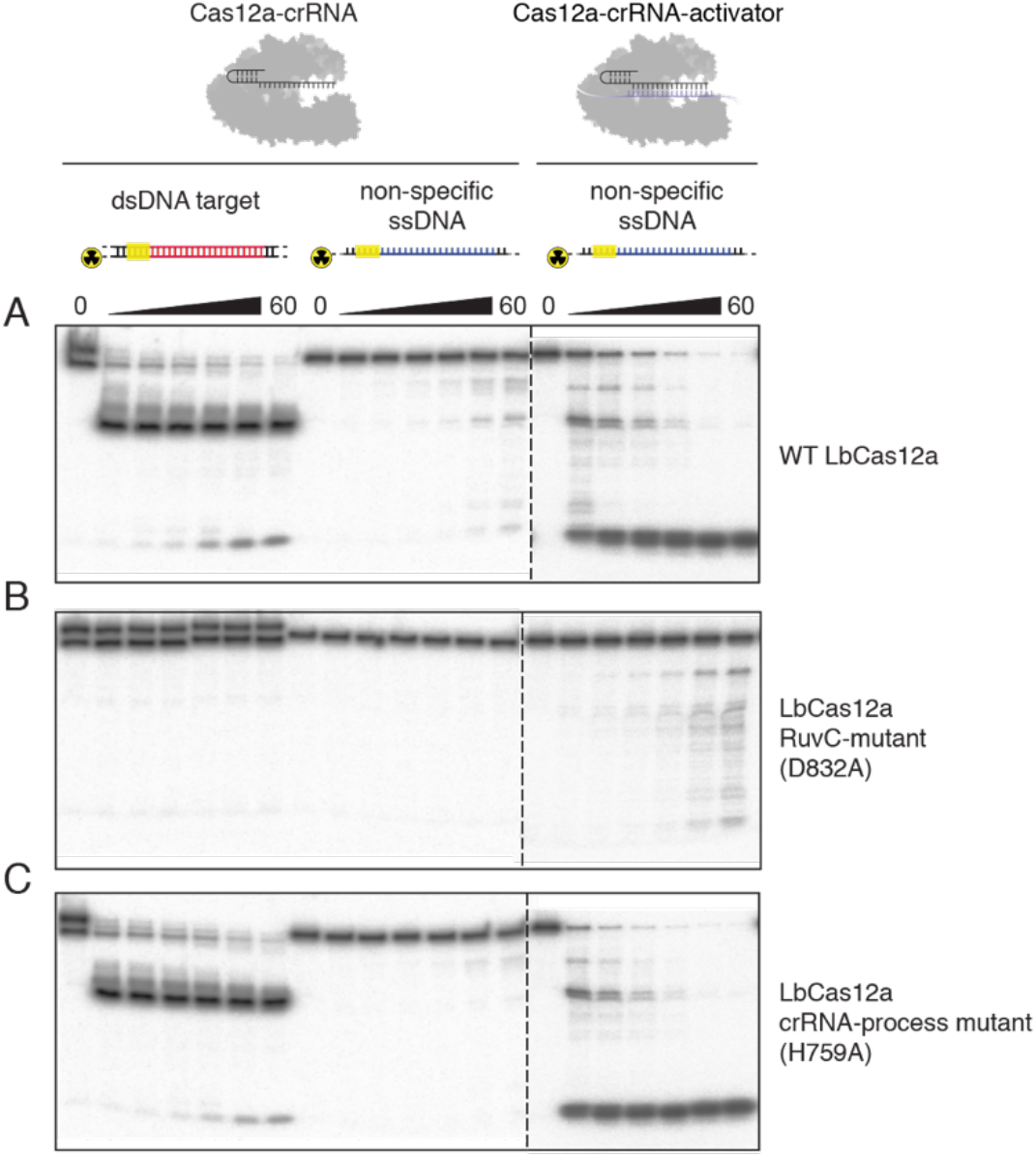
The RuvC nuclease is responsible for activator-dependent, non-specific DNase activity. Cleavage timecourse gel with radiolabeled target dsDNA and and non-specific ssDNA subsrates using (**A**) WT LbCas12a, (**B**) RuvC catalytic mutant (D832A) and (**C**) crRNA-processing mutant (H759A), with or without a ssDNA activator. Timecourses represent minutes and leavage products are resolved by denaturing PAGE.

**Fig. S5.**
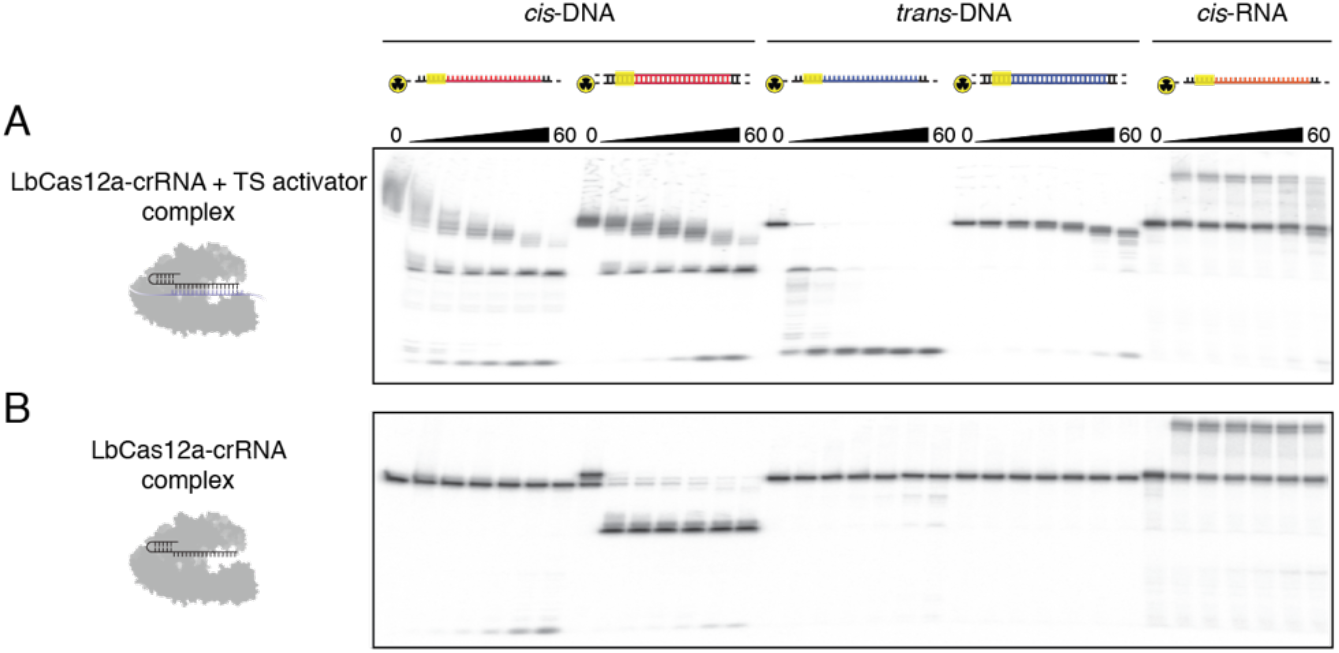
LbCas12a *trans*-cleavage degrades complementary and non-specific ssDNA, but not ssRNA. Cleavage timecourse gels of LbCas12a-crRNA complexes using (**A**) a ssDNA activator or (**B**) no activator with indicated radiolabeled substrates where *cis* indicates a complementary target and *trans* indicates a non-complementary sequence. Timecourses represent minutes and cleavage products are resolved by denaturing PAGE.

**Fig. S6.**
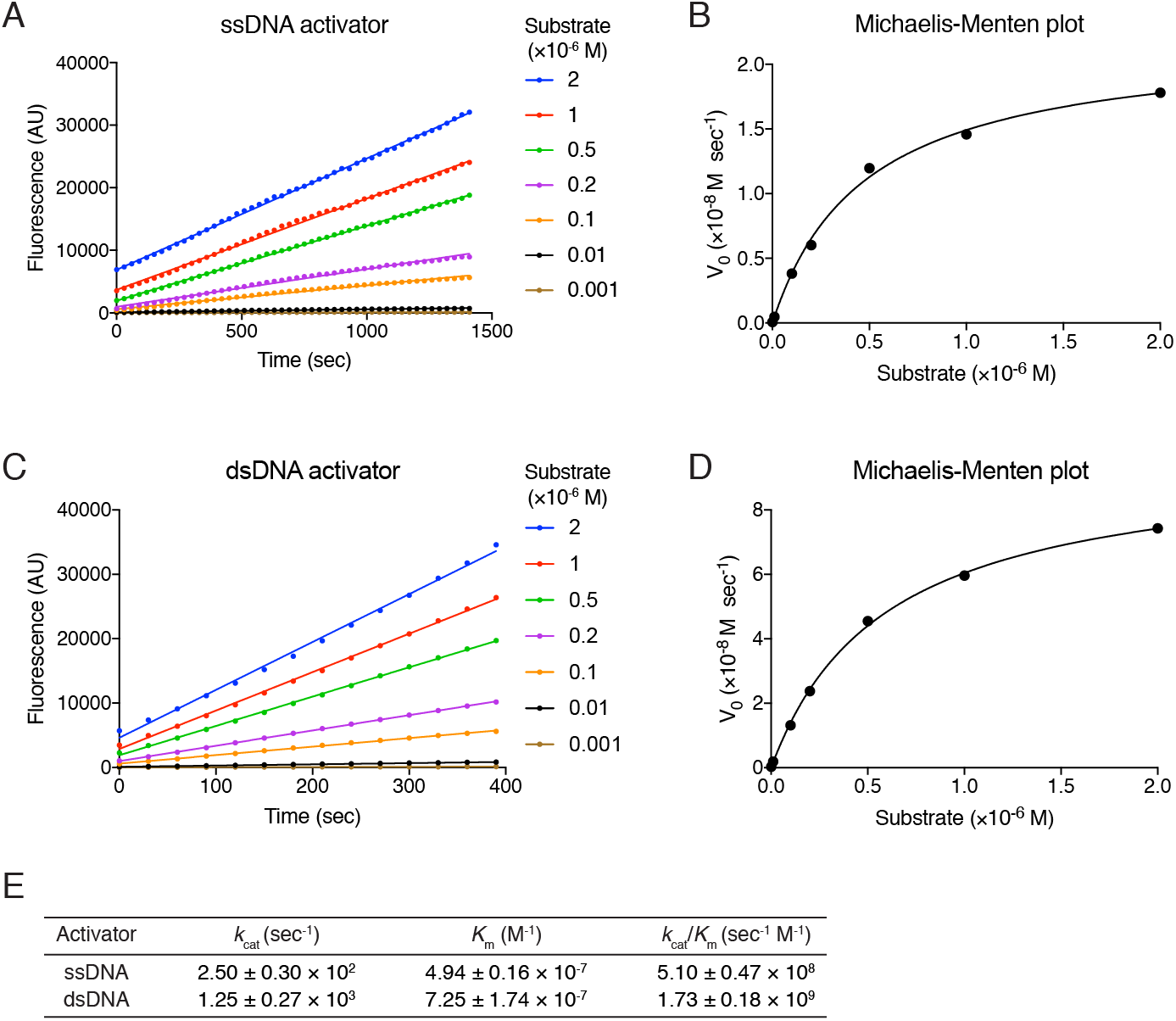
Michaelis-Menten analysis reveals robust *trans*-cleavage activity with a ssDNA and dsDNA activator. Representative plots of initial velocity versus time for a (**A**) ssDNA or (**C**) dsDNA activator, using 0.1 nM effective LbCas12a-crRNA-activator complex and increasing DNaseAlert substrate concentrations at 37°C. Michaelis-Menten fits for the corresponding (**B**) ssDNA or (**D**) dsDNA activator. (**E**) Calculated *k*_cat_, *K*_m_ and *k*_cat_/*K*_m_ values report the mean ± s.d., where *n* = 3.

**Fig. S7.**
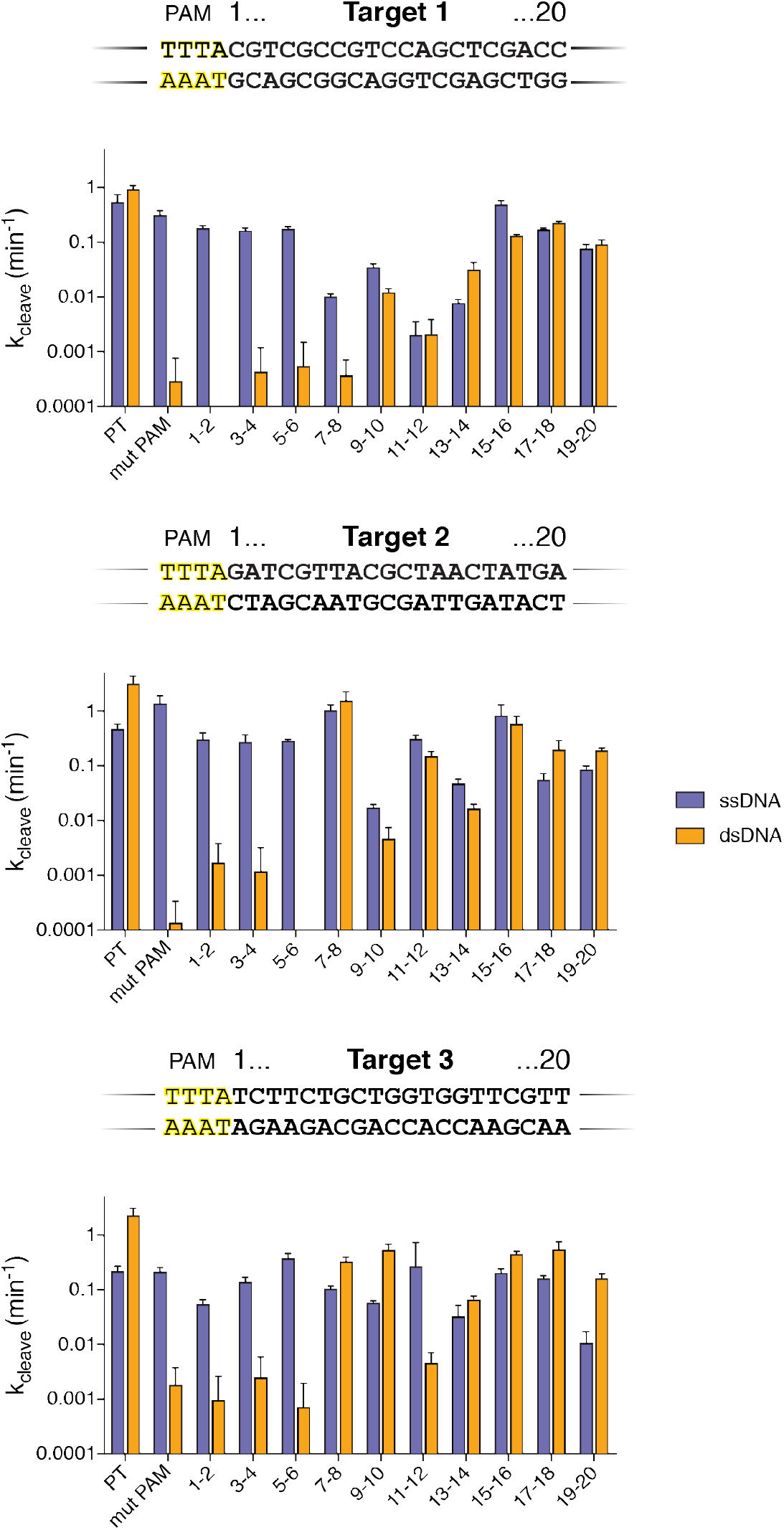
The PAM sequence and PAM-proximal mismatches in a dsDNA activator provide specificity for *trans*-activation. Quantification of *trans-cleavage* kinetics using mismatched substrates for three distinct target sequences; error bars represent the mean ± s.d., where *n* = 3.

**Fig. S8.**
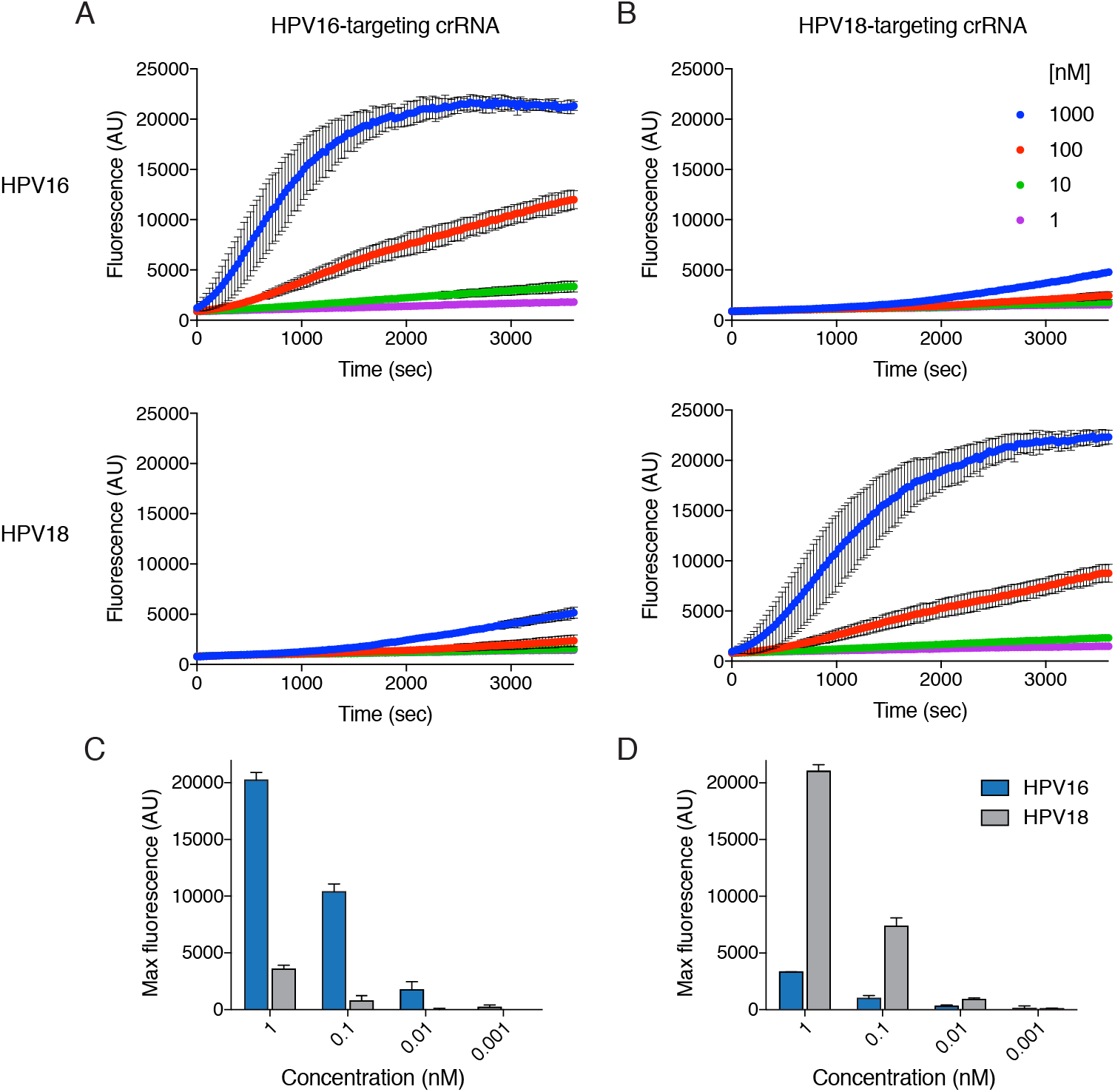
Human papillomavirus (HPV) detection assay timecourse. Fluorescence timecourses with LbCas12a preassembled with a crRNA targeting (**A**) HPV16 or (**B**) HPV16 in the presence of a dsDNA plasmid containing an HPV16 (top row) or HPV18 (middle row) genomic fragment and DNaseAlert substrate, with fluorescence measurements taken every 30 seconds for 1h at 37°C. (**C**) Maximum fluorescence signal obtained from timecourses in (**A**) and (**B**). Error bars represent mean ± s.d., where *n* = 3.

**Table S1.**
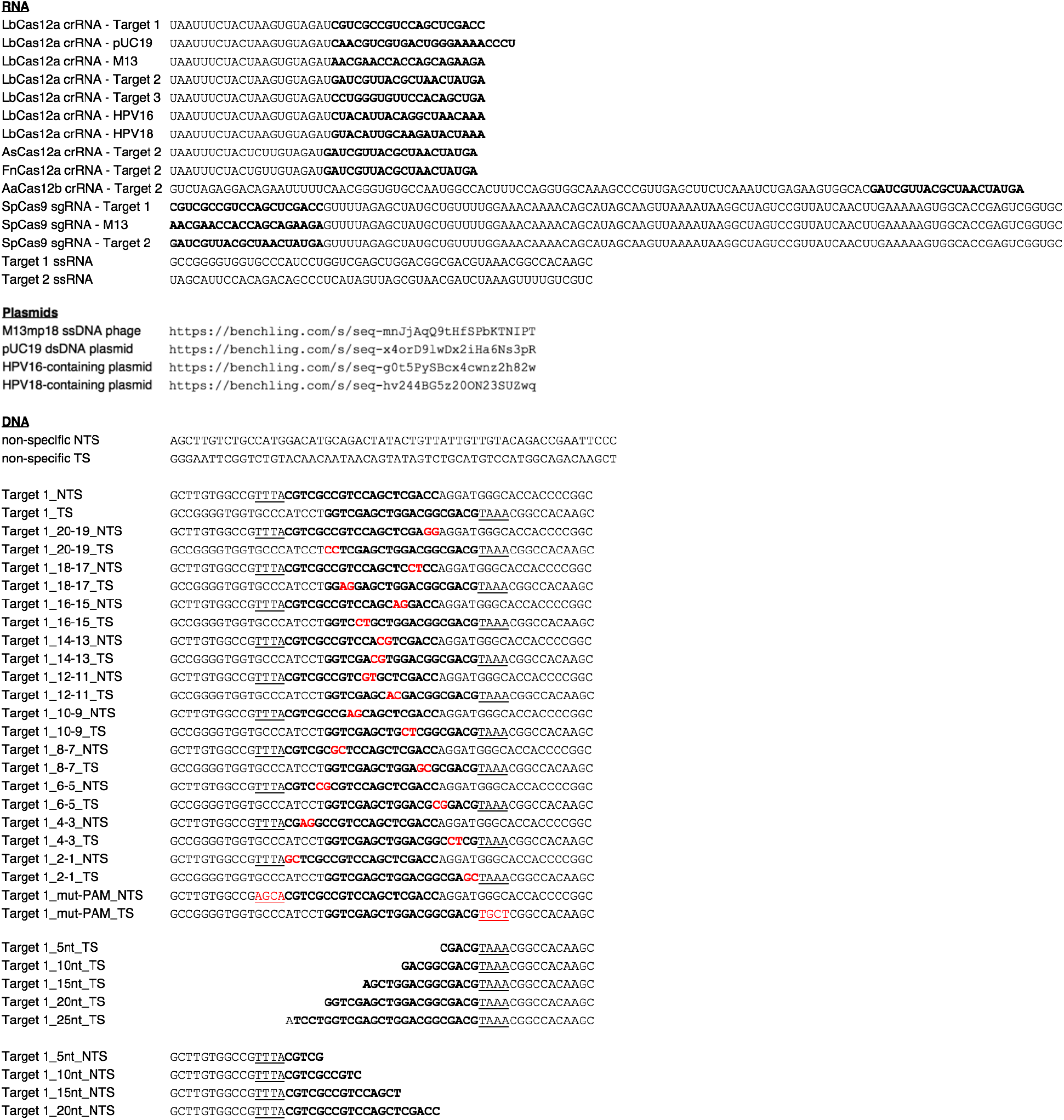
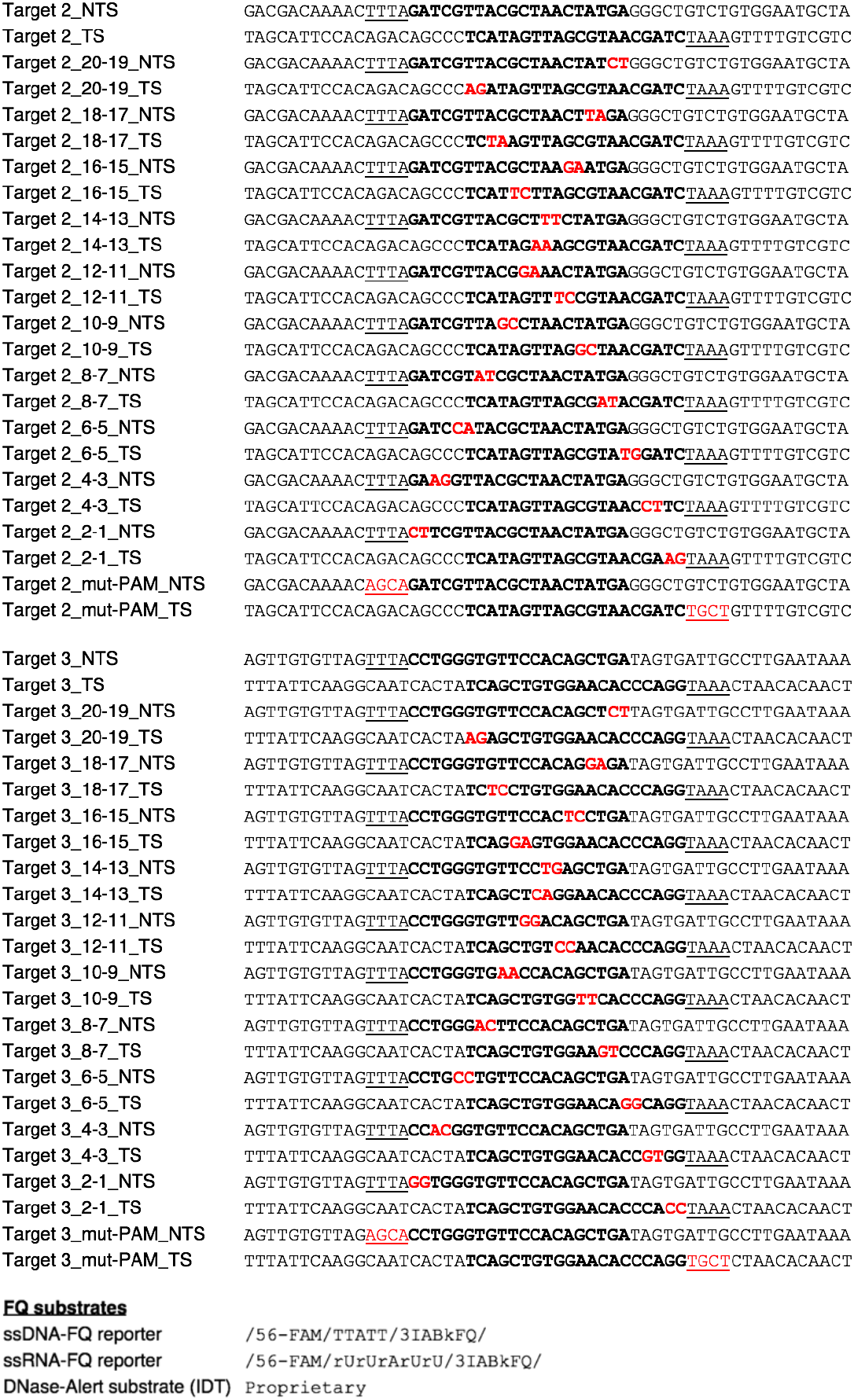
Nucleic acids used in this study.

